# Sperm AluY Epimutations in Male Partners of Couples with Unexplained Recurrent Pregnancy Loss

**DOI:** 10.64898/2026.09.24.753615

**Authors:** Elango Kamaraj, Jordan Moore, Mykle Keni, Anna Terry, Nastaran Salehisedeh, Neha Biju, Martha Susiarjo, Jonathon Hill, Timothy Jenkins, Winifred Mak

## Abstract

Recurrent pregnancy loss (RPL) affects 5% of couples worldwide. Over half of cases remain unexplained (uRPL) as paternal factors are rarely clinically evaluated or studied. This study presents data showing that sperm DNA methylation is a putative contributor to RPL. Using ONT direct whole methylome sequencing, we generated the largest cohort of locus-resolved, single-molecule sperm methylome maps from men with uRPL. We identified 294 differentially methylated regions (DMRs), of which 158 (53.7%) overlapped AluY repetitive elements, representing significant enrichment (z=40.9); most were hypomethylated in uRPL sperm and spanned the full length of individual AluY insertions. Because of single-molecule resolution, we show that the methylation differences found in uRPL sperm occur in a subpopulation of sperm rather than a uniform DNA methylation change across all sperm. AluY epimutation represents a previously unrecognized, predominant signature of paternal epigenetic perturbation in uRPL, establishing repetitive elements as a new axis of sperm epigenetic risk.

## Introduction

Recurrent pregnancy loss (RPL), defined as the loss of two or more pregnancies before 22 weeks of gestation^1^, affects approximately 5% of couples globally and represents a significant reproductive health burden, with psychological distress compounding the clinical complexity of each successive loss^2,3^. Decades of investigation have predominantly focused on maternal factors such as antiphospholipid syndrome, uterine structural anomalies, and endocrine dysfunction as contributing causes. Yet, these established categories collectively account for fewer than half of all RPL cases^4,5^. The remainder are classified as unexplained RPL (uRPL), leaving affected couples without a diagnosis, targeted treatment, or a clear path forward.

The paternal contribution to RPL has been understudied relative to maternal causes, with diagnostic evaluation of the male partner remaining minimal in routine clinical practice^6^. Emerging evidence implicates several paternal risk factors in RPL, including paternal age^7^, chromosomal translocations^8^, and sperm DNA fragmentation^9,10^. More recently, sperm epimutations have emerged as a particularly compelling mechanism for paternal contribution to RPL^11^, given that the spermatozoon contributes not merely a haploid genome but a structurally organized, epigenetically programmed nucleus whose chromatin architecture and methylation landscape are transmitted to the zygote and directly regulate early embryonic development.

Among these epigenetic marks, DNA methylation is the most chemically stable, germline-heritable, developmentally consequential in the mammalian genome, and essential for genomic imprinting, transposable element silencing, and early embryonic gene regulation^12,13^. A subset of sperm-derived methylation resists post-fertilization reprogramming and persists into the early embryo, directly influencing gene expression during zygotic genome activation^14,15^. Prior studies have begun to map this territory: aberrant methylation at imprinting control regions including *IGF2-H19*, *MEST*, *PEG3*, and *PEG10*, along with decreased global sperm 5-methylcytosine levels, have been reported in sperm from men with RPL^16^. More recent array-based and bisulfite sequencing studies have extended these findings to additional differentially expressed genes involved in embryonic development^11,17^. Together, these findings establish sperm epimutations as a biologically plausible and epidemiologically supported paternal risk factor for RPL. However, these prior studies are limited by either a targeted approach (arrays) or by very small sample sizes, such as a whole-genome bisulfite sequencing study that analyzed only 5 RPL and 5 control sperm samples^11^.

Furthermore, the contribution of epimutations as a mechanistic link to uRPL remains incomplete, as prior sperm methylome analyses have not adequately captured the repetitive elements. Repetitive elements, including SINEs and LINE-1 retrotransposons, constitute nearly half the human genome and are maintained in a densely methylated, silenced state throughout spermatogenesis by the PIWI-interacting RNA pathway^18^. Loss of repetitive element repression drives insertional mutagenesis, aberrant transcription factor binding, and dysregulation of neighboring genes that directly modulate germline integrity and embryonic gene regulation^19,20^. Repetitive regions have remained largely inaccessible to bisulfite-based DNA methylation methods, which, by design, exclude repetitive sequences because they cannot resolve individual repeat insertions with short reads^21^. Because repetitive element hypomethylation in sperm has been independently associated with unexplained male infertility and ART outcome, and LINE-1 hypomethylation has been reported directly in RPL-affected men^16,22,23^, there is a need to utilize technologies that enable analysis of repetitive sequences.

Oxford Nanopore Technology (ONT) long-read sequencing addresses this gap. By detecting 5-methylcytosine (5mC) and 5-hydroxymethylcytosine (5hmC) from the ionic current signal of native, unconverted DNA, ONT eliminates bisulfite-induced degradation and chemical ambiguity, while reads spanning tens of kilobases traverse full-length transposable element insertions and assign methylation status to individual locus-specific copies, a resolution inaccessible to short-read platforms^21,24,25^. Whole genome coverage without locus pre-selection enables unbiased, discovery-driven methylome analysis.

Here, we report ONT-based whole-genome sperm methylation profiling in 39 men from uRPL-affected couples and 30 fertile controls, providing the largest dataset of locus-resolved, single-molecule map of the sperm epimutation landscape across the complete methylome, and discovering a recurrent epimutation in the AluY repetitive elements as a candidate paternal determinant of recurrent pregnancy loss.

## Results

### Study Participants

A total of 69 men were enrolled in this study, comprising 39 male partners from couples with unexplained recurrent pregnancy loss (uRPL) and 30 fertile donor controls with at least one prior live birth. The uRPL cohort had experienced a median of 4 prior pregnancy losses (IQR 2 to 6), with the distribution spanning from 2 losses (38%) to 7 losses (2.6%). Of the 39 uRPL patients, 54% had no prior live births, 38% had one prior live birth, and the remaining 8% had two or three prior live births. The two cohorts differed significantly in age (p < 0.001), with a mean age of 32.18 ± 2.3 years in fertile donors and 35.45 ± 4.4 years in uRPL patients; paternal age was therefore included as a covariate in all downstream epigenetic analyses. Racial composition also differed between groups (p = 0.038). BMI was modestly higher in the uRPL group (median 26.40, IQR 24.95 to 28.55) compared to donors (median 24.90, IQR 23.70 to 27.05; p = 0.039), while smoking prevalence was comparable (p > 0.9). Sperm concentration (median 78 million per mL, IQR 49 to 129 versus 107, IQR 88 to 137; p = 0.025) and sperm motility (median 55%, IQR 45 to 73 versus 69%, IQR 60 to 78; p = 0.022) were significantly lower in uRPL patients though still within the normal limits for semen parameters per the 5th edition of the World Health Organization (WHO) laboratory manual for human semen examination. Participant characteristics are summarized in **Table 1**.

**Table 1.** Demographic, reproductive history, and semen characteristics of male partners from uRPL-affected couples and fertile donor controls.

| **Characteristic** | **Overall**<br>**n=69<sup>1</sup>** | **Donors**<br>**n=30<sup>1</sup>** | **Patient**<br>**n=39<sup>1</sup>** | **p-**<br>**value<sup>2</sup>** |
| --- | --- | --- | --- | --- |
| **Age** |  |  |  | <0.001 |
| 20-35 | 42 (61%) | 25 (83%) | 17 (44%) |  |
| 35-45 | 27 (39%) | 5 (17%) | 22 (56%) |  |
| **Race** |  |  |  | 0.038 |
| White | 56 (81%) | 21 (70%) | 35 (90%) |  |
| Other | 13 (19%) | 9 (30%) | 4 (10%) |  |
| **BMI** | 25.80 (24.40,<br>28.10) | 24.90 (23.70,<br>27.05) | 26.40 (24.95,<br>28.55) | 0.039 |
| **Current Smoker** |  |  |  | >0.9 |
| N | 65 (94%) | 28 (93%) | 37 (95%) |  |
| Y | 4 (5.8%) | 2 (6.7%) | 2 (5.1%) |  |
| **Number of**<br>**Miscarriages** |  |  |  |  |
| 2 | 15 (38%) | ND | 15 (38%) |  |
| 3 | 6 (15%) | ND | 6 (15%) |  |
| 4 | 10 (26%) | ND | 10 (26%) |  |
| 5 | 4 (10%) | ND | 4 (10%) |  |
| 6 | 3 (7.7%) | ND | 3 (7.7%) |  |
| 7 | 1 (2.6%) | ND | 1 (2.6%) |  |
| **Live Birth** |  |  |  |  |
| 0 | 21 (54%) | ND | 21 (54%) |  |
| 1 | 15 (38%) | ND | 15 (38%) |  |
| 2 | 2 (5.1%) | ND | 2 (5.1%) |  |
| 3 | 1 (2.6%) | ND | 1 (2.6%) |  |
| **Sperm Count** | 96 (72, 134) | 107 (88, 137) | 78 (49, 129) | 0.025 |
| **Sperm Motility** | 60 (53, 76) | 69 (60, 78) | 55 (45, 73) | 0.022 |
| <sup>1</sup>n (%); Median (IQR); no data (ND) |  |  |  |  |
| <sup>2</sup>Pearson's Chi-squared test; Wilcoxon rank sum test; Fisher's exact test |  |  |  |  |

### Global DNA Methylation in Sperm is Comparable between uRPL and Fertile Controls

Direct whole-genome methylation profiling was performed on all 69 samples using the Oxford Nanopore Technologies Native Barcoding Kit (NBK) for library preparation and sequencing. Filtered reads were aligned to the human reference genome (hg38). The fraction of CpG sites covered was 0.97 in fertile controls and 0.96 in uRPL patients (**Figure 1a**), and the median number of reads per site was 5.64× in controls and 5.96× in uRPL patients (**Figure 1b**), indicating near-complete CpG coverage at comparable depth (p=0.49) across groups. Promoter methylation levels followed a bimodal distribution as expected, with a high fraction of promoters showing very low methylation (0-10%) and the remainder predominantly methylated above 70% (**Figure 1c**), consistent with established patterns in human sperm^26^. Genome-wide methylation across non-overlapping 1 kb bins showed that most genomic regions were highly methylated (>70%; **Figure 1d**). Mean methylation was 0.752 among controls and 0.750 in uRPL patients while the median was 0.753 in controls and 0.751 in uRPL patients (**Figure 1e**), with no significant difference (p=0.26) between groups. Principal component analysis (PCA) of mean methylation levels in 10 kb genomic intervals across all 69 samples revealed no technical outliers, and samples did not separate by group (PC1 = 8.09%, PC2 = 5.82%; **Figure 1f**), confirming that genome-wide methylation profiles are comparable (Permanova, p=0.37) between uRPL and fertile sperm.

**Figure 1:**
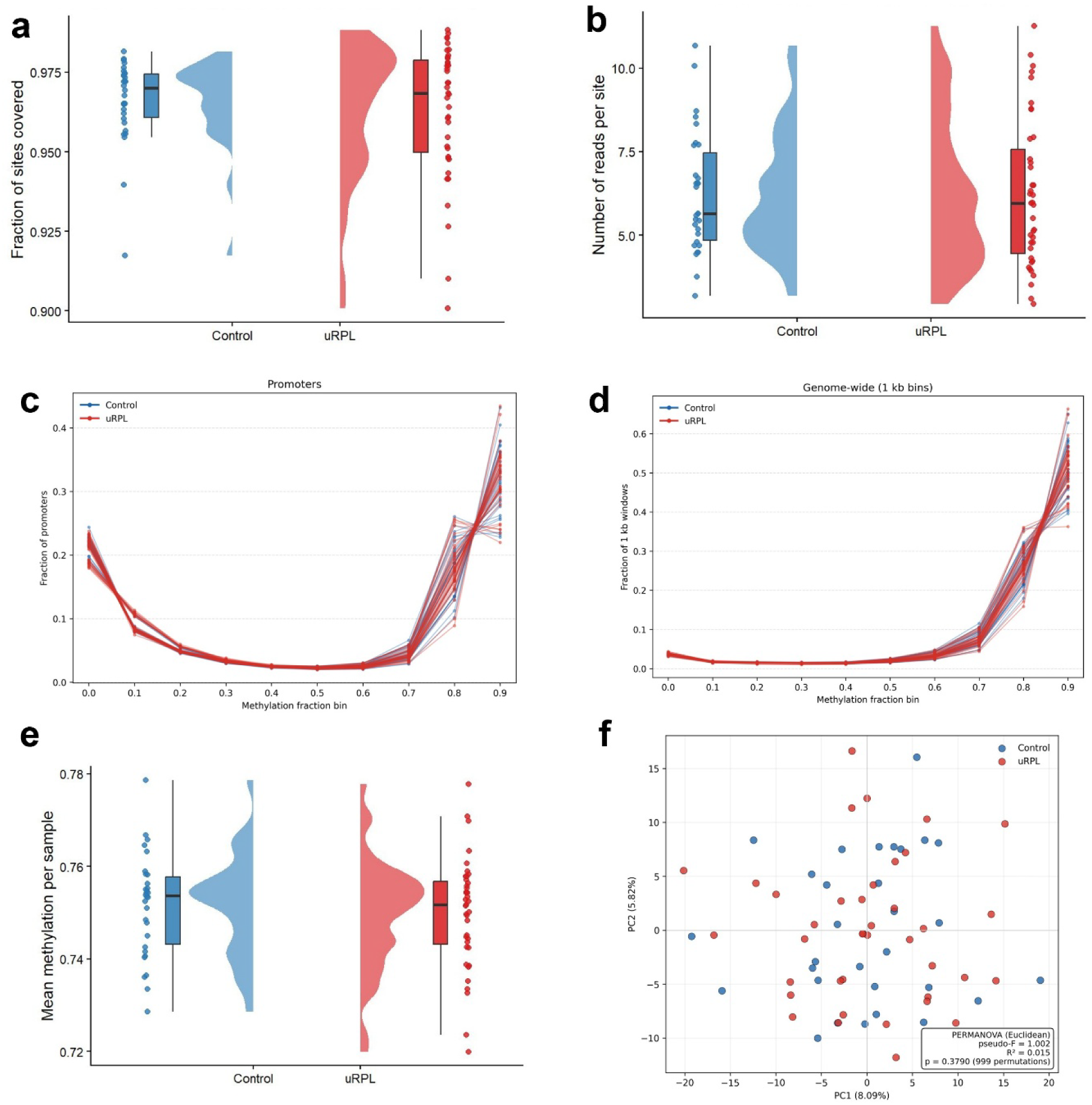
Comparable sequencing quality and global sperm methylome profiles between fertile and uRPL donors. **a**, Fraction of CpG sites covered by sequencing reads per sample, shown separately for fertile donors and uRPL patients, showing no bias between groups. **b**, Sequencing depth of coverage (number of reads per CpG site) per sample, showing no bias between the groups. **c**, Distribution of mean methylation levels across gene promoters, showing the expected bimodal pattern of largely unmethylated (0-10%) and methylated (>70%) promoters. **d**, Distribution of mean methylation levels across non-overlapping 1 kb genomic bins genome-wide. **e**, Mean genome-wide methylation level per sample by group; no significant difference was observed between groups. **f**, Principal component analysis (PCA) of mean methylation levels across 10 kb genomic intervals for all 69 samples, colored by group; samples did not separate by group (Permanova p= 0.37), and no technical outliers were identified. n = 30 fertile donors, 39 uRPL patients.

### Differentially Methylated Regions (DMRs) are Predominantly Hypomethylated and Enriched in Repetitive Elements

DMRs were identified using the combined 5-methylcytosine and 5-hydroxymethylcytosine modification signal from ONT sequencing and were defined as genomic regions containing at least five consecutive CpG sites that were within 1 kb of one another, showing differential modification in the same direction, and were significantly differentially modified between groups (p < 0.01). This analysis identified 316 DMRs, of which 22 were excluded following adjustment for paternal age; further adjustment for BMI and race did not exclude any DMRs. Therefore, 294 DMRs (**Supplementary Data. 1**) distinguished uRPL from fertile donor sperm (mean length 815 bp; median 476 bp). When we analyzed 5-hydroxymethylcytosine independently, we found no differentially hydroxymethylated sites, indicating that the detected methylation differences reflect alterations in 5-methylcytosine specifically. A volcano plot demonstrated that most DMRs were hypomethylated in uRPL sperm (256/294, 87.1%), whereas only 38 (12.9%) were hypermethylated (**Figure 2a**). The mean methylation difference was -11.4 percentage points (range: -1.1% to -21.2%) among hypomethylated DMRs and +8.5 percentage points (range: 1.0% to 18.0%) among hypermethylated DMRs. PCA restricted to DMR loci separated most uRPL patients from fertile donors (Permanova, p=0.001; **Figure 2b**), whereas genome-wide methylation profiles showed substantial overlap between groups. Genomic annotation revealed 161 intergenic (54.8%), 107 intronic (36.4%), 10 exonic (3.4%), and 16 promoter (5.4%) DMRs (**Figure 2c**). Strikingly, 276 of 294 DMRs (93.9%) overlapped repetitive-element regions, pointing to the repetitive-element as the primary site of sperm epigenetic changes in uRPL.

**Figure 2:**
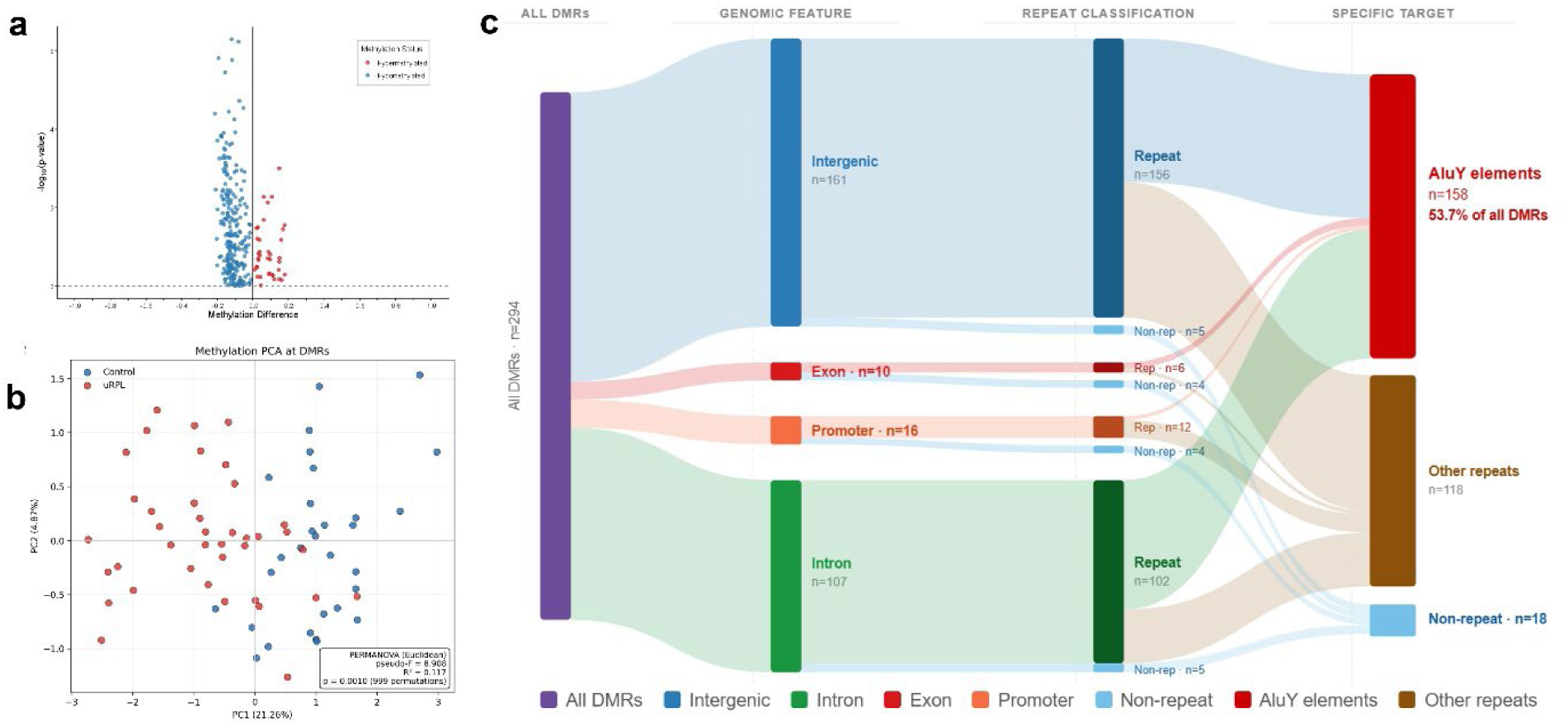
Differentially methylated regions distinguish uRPL from fertile sperm. **a**, Volcano plot of DMRs, showing methylation difference (uRPL–fertile donors) on the x-axis and statistical significance ([−log10 p-value]) on the y-axis. DMRs are highlighted, colored by direction of change (256 hypomethylated, 38 hypermethylated, out of 294 total DMRs). **b**, Principal component analysis (PCA) of methylation levels restricted to the 294 DMR loci, colored by group; unlike the global methylation profile (Figure 1f), uRPL patients largely cluster separately from fertile donors (Permanova p = 0.001). **c**, Genomic annotation of the 294 DMRs by feature type (intergenic, intronic, exonic, promoter) and repeat classification.

### AluY Elements are the Primary Targets of Differential Methylation in uRPL Sperm

Annotation of the 294 DMRs revealed an enrichment overlapping AluY elements, the evolutionarily youngest and transcriptionally competent subfamily of Alu SINEs^27,28^. Of 294 DMRs, 158 (53.7%) overlapped AluY elements, despite AluY comprising only 1% of the human genome across approximately 150,000 individual insertions. The expected number of overlaps by chance is 13.1, making the observed 158 overlaps approximately 40.9 standard deviations above expectation (z = 40.9), confirming that AluY enrichment is not attributable to random genomic distribution. Hypomethylation was distributed coordinately across all 21 CpG positions within the AluY consensus sequence (**Figure 3a**), indicating element-wide epigenetic aberration rather than focal disruption at specific CpGs. The majority of DMRs spanned the central and distal body of the AluY element, collectively covering its full 300 bp length (**Figure 3b**). The median DMR size of 304 bp was concordant with the median AluY insertion length of 305 bp (**Figure 3c**), confirming that many AluY DMRs correspond to the complete span of individual retroelement insertions rather than flanking sequences. As a representative example, the only DMR at the Sperm Associated Antigen 17 (SPAG17) locus, located within a single AluY insertion (**Figure 3d**), illustrates how AluY hypomethylation can constitute the sole epigenetic alteration at a functionally relevant sperm-expressed gene. To our knowledge, this is the first report of AluY hypomethylation as a shared feature of sperm epimutation in men with uRPL.

**Figure 3:**
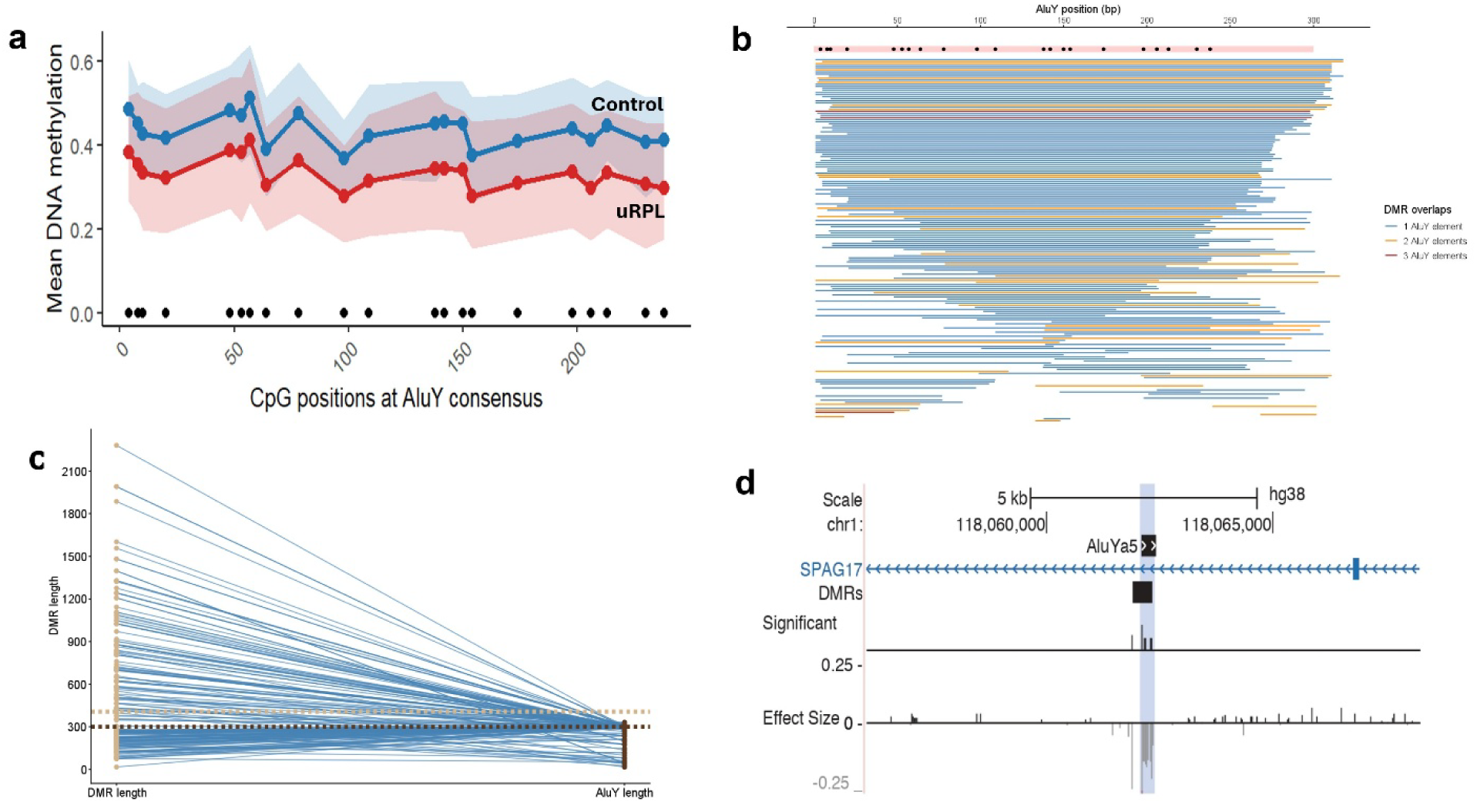
AluY repetitive elements are the predominant site of DMR enrichment. **a**, Methylation difference (uRPL - fertile) at each of the 21 CpG positions within the AluY consensus sequence, showing coordinated hypomethylation across the full element rather than focal disruption at individual or few CpGs. **b**, Positional distribution of AluY DMR overlaps along the AluY consensus sequence, showing that the majority of DMRs span the central and distal body of the element, collectively covering its full 300 bp length. **c**, Distribution of DMR sizes compared with the distribution of AluY insertion lengths genome-wide, showing concordance between median DMR size (304 bp) and median AluY insertion length (305 bp). **d**, Representative genome browser view of the single DMR at the Sperm Associated Antigen 17 (SPAG17) locus, located at an AluY insertion.

To independently validate AluY hypomethylation identified by ONT sequencing, we performed targeted bisulfite pyrosequencing at a representative Chr1 AluY DMR spanning an entire AluY element of 311 bp (chr1:161235257-161235567) across all study samples. Two sequencing regions were assayed: a 5-CpG region (chr1:161235409-161235461) and an adjacent 4-CpG region (chr1:161235547-161235568) covering 9 CpG sites in total. Consistent with the ONT findings, mean methylation across all 9 CpG sites was significantly lower in uRPL sperm compared to fertile donors (p = 0.03; **Supplementary Figure 1**). The magnitude of hypomethylation was comparable between methods: 17.7% by ONT and 14.58% by pyrosequencing, therefore independently confirming directional AluY hypomethylation at this locus and validating the quantitative accuracy of the ONT methylation calls. Pyrosequencing at the H19 ICR (chr11:1999924-1999996; 7 CpG sites) revealed no significant difference in mean methylation or at individual CpG sites between groups (p = 0.13), concordant with the ONT finding, confirming the absence of H19 ICR hypomethylation in this uRPL. Likewise, genome-wide LINE1 methylation didn’t differ significantly between groups (p=0.8). All three results are concordant with the corresponding ONT-based findings at these loci.

### uRPL-associated DMRs are Overrepresented in H2A.Z and Independent of Common Genetic Variation

To further characterize the genomic features of the uRPL sperm DMRs, enrichment analysis was performed for SNPs, histone modifications, and transcription factor binding motifs. DMRs showed significantly reduced overlap with common SNPs (odds ratio 0.17, q < 0.001), indicating that the methylation differences are not driven by known genetic sequence variants and represent genuine epigenetic alterations. DMRs showed significant underrepresentation in annotated H3K4me3 (odds ratio 0.29, q < 0.001), H3.3 (odds ratio 0.14, q < 0.001), H3K27me3 (odds ratio 0.62, q < 0.01), and bivalent chromatin domains (odds ratio 0.35, q < 0.01; **Figure 4a**). In contrast, DMRs were significantly overrepresented for the histone variant H2A.Z (odds ratio 15.8, q < 0.01), which marks nucleosome-retained regions within the otherwise protamine-packaged sperm chromatin^29^. No significant overrepresentation was identified for any of 472 known TF binding motifs (**Supplementary Data. 2**), and 16 de novo candidate motifs were all flagged as false positives attributable to the conserved internal sequence structure of AluY elements, indicating that AluY hypomethylation does not generate preferential binding sites for known transcription factors relative to other hypomethylated regions in sperm.

**Figure 4:**
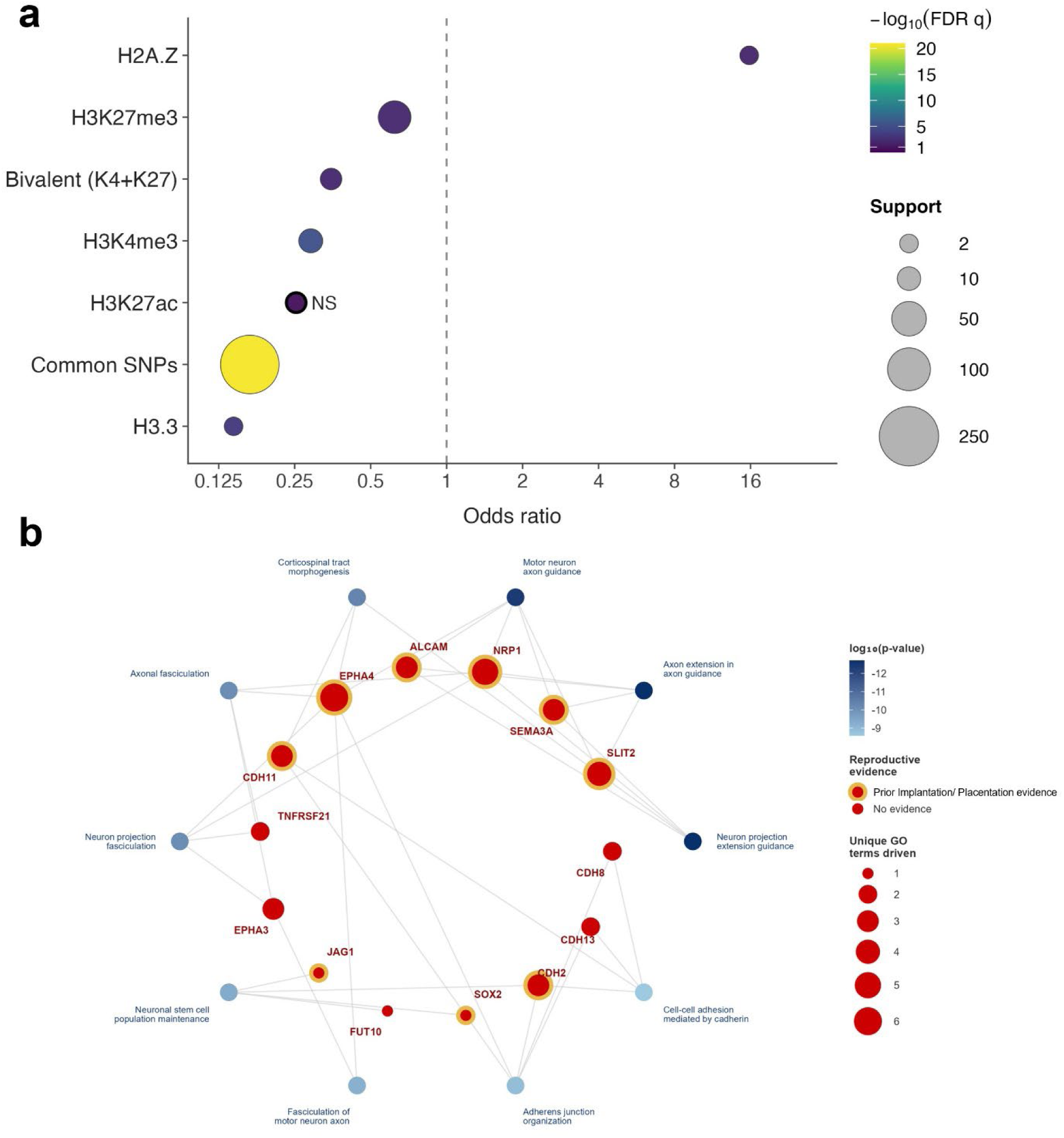
Chromatin context and gene ontology of DMR-associated genes. **a**, Overrepresentation (odds ratio, log scale) of DMRs for H2A.Z while underrepresented for common SNPs, H3K4me3, H3.3, H3K27me3, and bivalent chromatin domains, relative to genomic background. Odds ratios <1 indicate underrepresentation; >1 indicate overrepresentation. NS indicates non significance **b**, Ten most significantly enriched biological process terms among DMR-associated genes, with driving genes labeled for each term. Genes with previously reported roles in implantation and/or placentation are marked with an outer ring around their node.

### uRPL-associated DMRs are Associated with Genes Involved in Neuronal and Early Embryonic Development

Gene ontology analysis of DMR-associated genes identified 549 significantly enriched biological process terms (p < 0.05; **Supplementary Data. 3**). The most significantly enriched terms clustered around axon guidance and neuronal morphogenesis, including neuron projection extension involved in axon guidance (p = 2.99 × 10⁻⁶), motor neuron axon guidance (p = 4.22 × 10⁻⁶), and corticospinal tract morphogenesis (p = 3.32 × 10⁻⁵), driven by SLIT2, SEMA3A, NRP1, ALCAM, EPHA3, and EPHA4. Additional enriched clusters included neuronal stem cell population maintenance driven by JAG1, SOX2, and CDH2 (p=8.71×10⁻⁵); cadherin-mediated cell-cell adhesion (p = 1.87×10⁻^4^) and adherens junction organization (p = 1.37×10⁻^4^); driven by CDH2, CDH8, CDH11, and CDH13; and axonal fasciculation driven by EPHA3, EPHA4, TNFRSF21, and NRP1 (p = 4.10 × 10⁻⁵). The 10 highly enriched biological process terms and associated genes are shown in **Figure 4b**, and their prior independent reported roles relevant to early embryo development are listed in **Supplementary Table 1**.

### Single-Molecule Methylation Analysis Reveals Consistent Epigenetic Subpopulation Shifts in uRPL Sperm

Because of the advantages of the long-read technology used, we examined read-level methylation to determine whether the differential methylation reflects changes occurring uniformly across the entire sperm population (homogeneity) or whether only a subpopulation of sperm had unmethylated DMRs, with the rest remaining fully methylated (heterogeneity).

Examination of read-level methylation fraction distributions across all 294 DMRs revealed a shift toward the unmethylated state in uRPL, with an increased fraction of reads in the 0-5% methylation bin and a corresponding reduction in fully methylated reads (96-100%; **Figure 5a**). This pattern was consistent across most of DMRs. At representative loci, the read-level methylation distribution at chrX:61078676-61079581 was predominantly concentrated near 0% methylation in uRPL patients, whereas control reads spanned a broad range of methylation levels. At chr10:70918300-70919092, controls retained a prominent fully methylated peak near 100% that was markedly diminished in uRPL (**Figure 5b**). Analysis of DMRs with the greatest and smallest fraction of reads in the 0-5% methylation bin confirmed the systematic nature of these subpopulation shifts: top-ranked DMRs showed a majority of patient reads with near-zero methylation, while controls retained substantial methylated populations, and bottom-ranked DMRs showed the opposite pattern, corresponding to the minority hypermethylated DMR set (**Figure 5c**). Intra- and inter-sample variances at DMRs were comparable between uRPL patients and fertile controls (**Supplementary Figure 2**), supporting the conclusion that the 294 DMRs reflect consistent, reproducible group-level methylation differences rather than artifacts of inter-individual variability.

**Figure 5:**
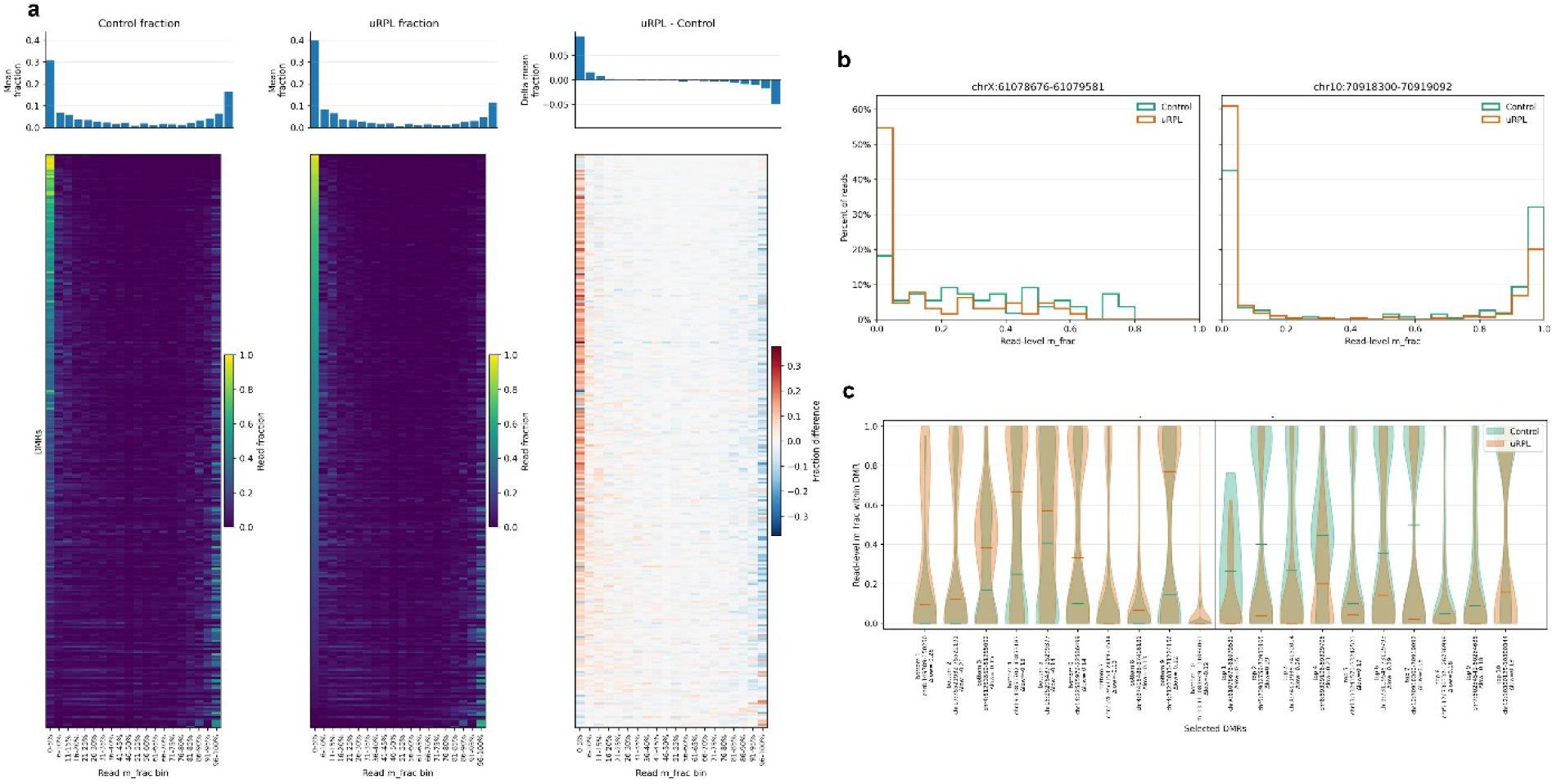
Single-molecule read-level analysis reveals a subpopulation shift. **a**, Distribution of read-level methylation fractions across all 294 DMRs, binned in 5% increments, comparing uRPL patients and fertile controls; uRPL sperm show an increased fraction of reads at 0-5% methylation and a reduced fraction at 96-100% methylation. (Rows correspond to DMRs, and the Columns correspond to methylation bins) **b**, Read-level methylation fraction distributions at two representative DMR loci. At chrX:61,078,676–61,079,581, patient reads are predominantly concentrated at 0% methylation while control reads span a broad distribution, illustrating complete subpopulation hypomethylation. At chr10:70,918,300–70,919,092, both groups show high concentrations at 0% methylation with a higher proportion of unmethylated reads in uRPL patients, while controls retain a substantially larger fully methylated peak near 100% compared to uRPL patients, illustrating both an enrichment of the unmethylated subpopulation and a marked reduction of the fully methylated subpopulation in uRPL sperm. **c**, Overlaid read-level methylation fraction distributions for the ten bottom-ranked (left panel) and ten top-ranked (right panel) DMRs based on smallest and greatest fraction of reads in the 0-5% methylation bin. Top-ranked DMRs show uRPL patient reads concentrated near 0% methylation while control reads span a broader distribution with higher median values, confirming locus-level hypomethylation in uRPL sperm. Bottom-ranked DMRs show the inverse pattern, with control reads concentrated near 0% and uRPL patient reads shifted toward higher methylation values, corresponding to the minority hypermethylated DMR set.

## Discussion

The present study provides the first locus-resolved, single-molecule characterization of sperm DNA methylation alterations associated with uRPL and represents the largest genome-wide sperm methylome analysis performed in a well-phenotyped cohort of male partners of uRPL couples. Our identification of AluY elements as the predominant genomic context of sperm epigenetic dysregulation reveals a previously unrecognized dimension of paternal epigenetic risk in uRPL, with AluY-associated DMRs enriched 40.9 standard deviations above the expected genomic distribution. Our findings build upon previous observations linking repetitive element methylation with male reproductive failure. Urdinguio et al.^22^ found DNA methylation at repetitive sequences including AluYB8, to be significantly lower in spermatozoa from males with unexplained infertility than in fertile controls, suggesting that hypomethylation of repetitive elements is a broader feature of male reproductive failure. El Hajj et al.^23^ reported reduced Alu methylation in sperm samples associated with ART cycles resulting in miscarriage compared with those resulting in successful pregnancy. Together, these observations suggest that Alu hypomethylation may extend across different male reproductive failure phenotypes.

Previous studies of sperm methylation in uRPL relied on methylation arrays, targeted pyrosequencing of imprinted loci, or whole-genome bisulfite sequencing. These studies identified alterations at loci such as H19 ICR, but their findings have been inconsistent across cohorts^16,17,30^. A whole-genome methylation study expanded the search beyond imprinting regions^11^; however, the use of bisulfite-based methodologies limited resolution to population-averaged methylation estimates across DNA molecules. Although H19 ICR methylation has been repeatedly proposed as a candidate paternal epigenetic marker, we did not detect differential H19 methylation by either nanopore sequencing or targeted pyrosequencing in our cohort. This discrepancy may reflect differences in sample size, the definition of reproductive phenotype, or methodological approaches.

Furthermore, a systematic review and meta-analysis identified evidence of publication bias and considerable variability among studies evaluating H19 methylation in men with infertility or RPL^32^. Moreover, recent evidence suggests that some reported imprinting abnormalities in sperm may reflect somatic cell contamination rather than true sperm-specific epimutations^31^. Somatic contamination is unlikely to explain the AluY signal reported here, as contamination by somatic Alu sequences would bias the signal toward hypermethylation, not the hypomethylation observed^33–35^. Collectively, our findings expand the current understanding of paternal epigenetic risk in uRPL by shifting the focus beyond imprinting loci and identifying AluY hypomethylation as a major, previously unrecognized signature of uRPL sperm.

The observed hypomethylation in the current study is concentrated within a single repeat subfamily, a pattern more consistent with a locus-selective surveillance defect rather than the global genome-wide failure of methylation seen with broad DNMT3A/DNMT3B loss^36^. The most established locus-selective silencing mechanism for transposons in the male germline is the PIWI-interacting RNA pathway, which specifically directs de novo methylation machinery to transposon loci, including AluY elements^18,19^, and inherited defects in the piRNA pathway are known to cause transposon derepression and infertility in humans^37^. Because AluY is the youngest and most recently active Alu subfamily, it likely receives disproportionately dense piRNA targeting compared with older, evolutionarily stable subfamilies, consistent with the established relationship between transposon age/activity and piRNA-mediated silencing in humans and other mammals^38^. This dependence on active, ongoing surveillance may make AluY a plausible weak point if that surveillance falters. We therefore propose deficient piRNA-directed surveillance of AluY elements as a candidate mechanism, and future studies should focus on piRNA sequencing and PIWI gene expression profiling in uRPL men.

Since AluY hypomethylation originates during spermatogenesis, its consequences could plausibly extend in two directions after fertilization: disrupted inheritance of sperm chromatin architecture and aberrant transposon promoter activity during early embryonic reprogramming, neither of which is mutually exclusive. We consider each in turn, followed by a third and independent consequence: genome instability within the sperm itself (**Supplementary Figure 3**).

We first consider chromatin inheritance. Unlike the imprinting defects reported in prior RPL studies, uRPL DMRs were significantly underrepresented in annotated H3K4me3, H3.3, H3K27me3, and bivalent chromatin domains, the very marks that characteristically define imprinting control regions in sperm. Instead, they are overrepresented in H2A.Z, a histone variant marking nucleosome-retained regions within the otherwise protamine-packaged sperm genome. H2A.Z enrichment has previously been reported to concentrate largely on pericentric heterochromatin rather than developmental gene loci^12^, making its strong enrichment at AluY-associated DMRs in our dataset a distinct and unexpected pattern. More broadly, the DNA methylation state of sperm is known to shape which genomic regions retain nucleosomes and which genes receive active histone marks after fertilization^39^. AluY hypomethylation at these H2A.Z-marked loci may therefore do more than alter a single molecular mark: as a structural, packaging-level effect, it could disrupt the chromatin architecture the embryo inherits from the sperm, with consequences for embryonic gene regulation that remain to be defined^40^.

A second, related mechanism involves Alu promoter activity itself. Alu elements contain an internal RNA Pol III promoter and are preferentially distributed in gene-rich regions, where they regulate neighboring gene expression through effects on transcription, alternative splicing, and RNA stability^41^. If AluY elements enter the embryo in a hypomethylated state from the paternal genome, their promoter activity may become aberrantly accessible during the programmed demethylation wave targeting Alu elements in preimplantation development, potentially disrupting the ordered gene expression program of zygotic genome activation^15,42^. This reasoning gives biological plausibility to the gene ontology enrichment pattern observed in our data: although the most significantly enriched biological process terms carry neuronal GO annotations, DMR-associated genes converge on processes critical to implantation, placentation, and early embryonic development (**Supplementary Table 1**), a pattern consistent with, though not direct evidence for, a Pol III-mediated dysregulation mechanism at these loci. Genes including SLIT2, NRP1, JAG1, and the EPHA4 receptor family have reported roles in trophoblast migration and placental vascularization^43–49^, while SEMA3A, ALCAM, SOX2, CDH2, and CDH11 contribute to blastocyst-endometrium interaction, trophectoderm formation, and decidualization^50–58^.

Independent of these embryo-facing consequences, AluY hypomethylation may also directly compromise the sperm genome. Young Alu elements including AluY, remain retrotranspositionally active and are prone to R-loop formation and DNA double-strand breaks (DSBs) in mature sperm^59–61^, and elevated sperm DSBs have independently been linked to recurrent pregnancy loss^10^. The zygote has a limited capacity to repair paternal DNA damage, and damage exceeding this threshold has been linked to pregnancy failure^62,63^. This raises the possibility that AluY hypomethylation contributes to sperm genome instability itself, independent of any effect on the embryo’s chromatin or transcriptional program. A related possibility is that AluY retrotransposition disrupts developmental genes through direct insertional mutagenesis, an established disease mechanism for Alu and other retrotransposons^64^. Insertions arising during spermatogenesis would be present throughout the fertilizing sperm genome as dominant mutations, whereas continued AluY transcriptional activity after fertilization could generate mosaic insertions with either dominant or recessive effects depending on the affected lineage. Testing this hypothesis directly will require the detection of de novo AluY insertions in sperm or in products -of- conception.

These mechanistic findings also carry clinical implications. Brogaard et al.^65^ reported that men with abnormal sperm epigenetic profiles have significantly lower pregnancy success rates despite similar sperm motility and concentration, underscoring the clinical potential of epigenetic sperm testing beyond conventional semen analysis. Our findings suggest that AluY methylation at representative loci could complement the diagnostic framework for uRPL by capturing the dominant repetitive-element dimension of sperm epigenetic dysregulation. These observations come with several limitations: a modest age difference between groups remains a possible contributing factor despite covariate modeling, and the extent to which AluY methylation is transmitted to and retained in the embryo through the post-fertilization reprogramming wave remains unknown.

The single-molecule resolution of ONT revealed that AluY hypomethylation reflects a shift in the balance between unmethylated and fully methylated sperm subpopulations rather than a uniform reduction in methylation. This subpopulation architecture indicates that individual sperm within a single ejaculate differ substantially in their AluY methylation state, reflecting epigenetic mosaicism that may contribute to variation in the molecular state of sperm available for fertilization across conception attempts within the same couple. Together with recent observations of ongoing AluY retrotransposition in human sperm^59^, these findings highlight AluY elements as dynamic genomic regions exhibiting heterogeneity at both epigenetic and genetic levels. This genetic dynamism does not, however, extend to common sequence variation at these loci as DMRs showed significantly reduced overlap with common SNPs, confirming that the methylation differences detected in the current study reflect genuine epigenetic alterations rather than artifacts of common genetic variation. Future single sperm epigenomic approaches will be required to determine whether hypomethylation across multiple AluY loci occurs in a coordinated manner within individual sperm cells.

In summary, this study provides the first locus-resolved, single-molecule map of sperm epimutations in uRPL and identifies AluY hypomethylation as a newly discovered, dominant, and reproducible paternal epigenetic signature. The findings reveal the repetitive elements as a major and overlooked dimension of sperm epigenetic dysregulation in uRPL, distinct from the imprinting defects described in prior studies. AluY hypomethylation co-occurs with altered H2A.Z-marked chromatin inheritance and a network of developmental genes linked to implantation and placentation, pointing toward a model in which paternal epigenetic dysregulation at repetitive loci contributes to early pregnancy failure. A candidate piRNA surveillance defect and AluY-associated genome instability offer plausible, literature-supported explanations for the origin and downstream consequences of this signature. These findings open new directions for understanding the biology of recurrent pregnancy loss, for developing epigenetic biomarkers of paternal reproductive risk, and for reconsidering the clinical evaluation of the male partner in couples with unexplained recurrent pregnancy loss.

## Methods

### Study Participants

In this case-control study, we enrolled 39 uRPL couples with two or more pregnancy losses. All participants were recruited from UT Health San Antonio, UT Health Austin, and the University of Michigan. This study was approved by single IRB overseen by the IRB at the University of Texas Health San Antonio under protocol number 20230315HU (May 2^nd^, 2023); all participants provided written informed consent. Couples were included only if the RPL workup was negative according to American Society for Reproductive Medicine (ASRM) guidelines. Couples were excluded if the female partner had an identified cause of RPL such as antiphospholipid syndrome, uterine anomalies, parental chromosomal abnormalities, or endocrine disorders or significant comorbidities, including uncontrolled diabetes, hypothyroidism or renal failure.

Inclusion criteria required maternal age between 18 and 40 years and paternal age between 18 and 45 years. Demographic information (age, body mass index, smoking status, alcohol use, and drug use) and medical history, were collected for both partners, the male partners completed the Charlson Comorbidity Index, and all data was recorded in a HIPAA-compliant REDCap database. Control sperm samples were obtained from 30 closely age-matched fertile donors with at least one documented live birth, sourced from the Fairfax and Seattle sperm cryobanks. Men in the uRPL group were asked to provide a semen sample, and routine semen analysis was carried out. Neat semen was aliquoted into cryovials and cryopreserved using a glycerol-based cryoprotectant according to standard sperm preparation protocols provided by the cryobanks, ensuring that case and donor samples were processed comparably and minimizing sperm processing as a potential confounder in downstream analyses. Baseline characteristics were summarized by group (uRPL patients vs. donors) using descriptive statistics. Continuous variables were presented as medians with interquartile ranges (IQR) and compared using the Wilcoxon rank-sum test. Categorical variables were presented as frequencies and percentages, with group differences assessed using Pearson’s chi-squared test or Fisher’s exact test where cell counts were small. All analyses were performed in R (v.4.3.2), with the significance threshold set at α=0.05.

### DNA Extraction

Genomic DNA was extracted from sperm samples obtained from 39 men with uRPL and 30 fertile donors using a protocol adapted from Jenkins *et al.*^66^ and the DNeasy Blood & Tissue Kit (Qiagen). Briefly, 1 × 10*^7^*spermatozoa per sample were washed in phosphate-buffered saline (PBS) by centrifugation at 10,000 × g for 10 min. Removal of somatic cell contamination was performed by resuspending the sperm pellet in 10 mL somatic cell lysis (SCL) buffer (10 mM Tris-HCl, 50 mM KCl, 2.5 mM MgCl₂, 4 mM DTT, 0.05% w/v SDS, 0.5% v/v Triton X-100; pH 7.4), mixed thoroughly, and incubated overnight. The following day, samples were mixed and centrifuged at 800 × g for 10 min. The supernatant was removed, leaving approximately 100 µL to resuspend the pellet. An equal volume (100 µL) of buffer X2 (20 mM Tris-HCl, pH 8.0; 20 mM EDTA; 200 mM NaCl; 4% SDS) was added, followed by 12 µL Proteinase K (Qiagen) and 8 µL of 1 M DTT. Samples were incubated at 56°C for 2 h in a thermomixer with agitation at 300 rpm. Subsequently, 200 µL Buffer AL and 200 µL 100% ethanol were added, mixed thoroughly, and the lysate was loaded onto a DNeasy Mini spin column. Columns were centrifuged at 8,000 × g for 1 min and transferred to new collection tubes. Wash steps were performed with 500 µL Buffer AW1 (8,000 × g for 1 min) followed by 500 µL Buffer AW2 (20,000 × g for 3 min). Columns were then transferred to clean 1.5 mL microcentrifuge tubes, ensuring no ethanol carryover. DNA was eluted by adding 60 µL Buffer AE directly to the membrane, followed by incubation at room temperature for 20 min and centrifugation at 8,000 × g for 1 min. A second elution was performed with 40 µL of Buffer AE. DNA concentration was measured using both NanoDrop spectrophotometry and Qubit fluorometry.

### DNA Library Preparation and Sequencing

Genomic DNA was mechanically sheared to an average fragment length of 10 kb and concentrated to a minimum input of 200 ng using a SpeedVac concentrator. Sequencing libraries were prepared using the Oxford Nanopore Technologies Native Barcoding protocol (SQK-NBD114.96) with minor volume adjustments. Barcoded libraries were pooled at three to four samples per flow cell and sequenced on PromethION Flow Cells (FLO-PRO114M) using a PromethION 2 instrument.

### Data preparation

MinKNOW was used to concurrently basecall POD5 files with dna_r10.4.1_e8.2_400bps_hac@v4.3.0 while sequencing. Basecalled reads were output in unaligned BAM files and subsequently aligned using dorado version 1.1.1 with the UCSC GRCh38 primary assembly (hg38)^67^. First, we used modkit (version 0.5.0)^68^ modbam adjust-mods to combine 5hmC probability scores with 5mC probability scores. We also used modkit call-mods to convert modification probability scores (ML), which is a linear scale (0-255), to a binary call (0 or 255). DNMTools (version 1.5.1) counts-nano was then used to summarize modification calls by genomic position, while modkit extract calls were used to extract individual, single-molecule modification calls for subsequent analyses. Additionally, sample global methylation was analyzed for potential somatic cell contamination, of which we found none. Genome-wide and promoter methylation was analyzed using PCA and PERMANOVA tests to determine any potential batch effects between samples sequenced.

### DMR Calling and Genomic Distribution

Using the output from DNMTools counts-nano, we utilized DNMTools radmeth to identify differentially methylated CpG sites (5mC + 5hmC). Radmeth^69^ employs a beta-binomial regression framework that accounts for variability beyond that expected under a binomial model and permits adjustment for clinical covariates. Using Radmeth’s results, we used a sliding-window approach to identify DMRs by merging at least five consecutive, significantly differentially methylated CpG sites that were within 1 kb of one another, showed methylation changes in the same direction (e.g., hypermethylation), and met a p-value cutoff of 0.01. The same process was repeated independently for age, race and BMI to assess their influence. No DMRs were associated with race and BMI, whereas 22 DMRs significantly associated with age (p < 0.05) were identified and subsequently removed from the uRPL-associated DMR set, yielding the final 294 DMRs used in all downstream analyses.

Genomic distribution of DMRs was analyzed by annotating DMR intervals with the hg38 assembly^67^. Specifically, gene annotations were assigned using the UCSC NCBI RefSeq annotation track, and repeat regions were assigned using the UCSC RepeatMasker annotation track^70,71^. The transcription start site (TSS) was defined as the transcript start coordinate on the sense strand and the transcript end coordinate on the antisense strand. Promoters were defined as regions spanning 2 kb upstream and 500 bp downstream of the TSS. DMRs were annotated as overlapping a promoter, exon, intron, intergenic region, repeat element, and/or AluY element if the DMR start position was less than the annotation stop position and the DMR end position was greater than the annotation start position. For AluY annotation, all AluY subfamily members were collectively annotated as AluY. Intervals were subsequently created for DMR regions that intersected AluY repeat annotations. For each sample, dnmtools counts and dnmtools selectsites were used to extract DNA methylation calls from its aligned BAM file within these intervals^72^. For each call within each interval, the genomic position was converted to a relative position inside the AluY consensus sequence^73^. For each relative CpG position in the AluY sequence, control and uRPL mean methylation fractions were calculated.

### Read-Level Methylation Distribution and Intra- and Inter-Sample Variance Analysis

Modkit extract calls-generated parquet files (ch3 format)^74^ was filtered for 5mC modifications at CpG positions whose start position were located within DMR intervals. Read 5mC fraction was calculated by dividing total 5mC calls by the total number of calls at positions within the DMR. The variance between all reads at each given DMR constituted a distribution from which intra-sample variance was estimated. Likewise, the average read 5mC fraction at a given DMR was used to determine variance between samples for all DMRs, from which distribution we estimated inter-sample variance. This was repeated for both fertile donor controls and uRPL samples.

### SNPs, Histone Marks, and Transcription Factor Binding Enrichment analysis

Hypomethylated regions (HMRs) in the sperm methylome that did not intersect with DMRs were used as the background model for SNP and histone mark enrichment analyses. SNP annotations were obtained from dbSNP build 155 mapped to hg38^67,75^. Sperm chromatin features, including H3K4me3, H3K27me3, H3K27ac, H3.3, H2A.Z, and a derived bivalent H3K4me3/H3K27me3 category, were obtained from publicly available sperm ChIP-seq datasets deposited under GEO accessions GSE15690 and GSE57095^12,76^. UCSC’s hg19ToHg38.over.chain.gz was used for the histone mark’s coordinate liftover^77^. LOLA, GenomicRanges, rtracklayer, regioneR, and dplyr were used to determine overlaps of SNPs and histone marks for each DMR and background HMR^78–84^. DMR intervals were randomly resampled using resampleRegions and 10,000 permutations were run for histone mark overlaps. For each feature, a two-sided Fisher’s exact test was run, comparing the number of DMRs with/without overlaps of said feature versus the number of background HMRs with/without overlaps of said feature. A Benjamini-Hochberg correction was applied, and significant enrichment and/or depletion deemed significant when q < 0.05.

For transcription factor (TF) analysis, HMRs lacking DMRs, identified from the same samples, were also used as the background model to control for sequence composition characteristics of HMRs and isolate enrichment specific to differentially methylated loci. All analyses were performed against the human reference genome hg38^67^. Known and de novo transcription factor (TF) motif enrichment were assessed on the DMR dataset using HOMER’s findMotifsGenome.pl (v5.1)^85^. Prior to enrichment testing, HOMER normalized background regions to match the GC content distribution of target DMRs. Significance was determined by binomial test with Benjamini-Hochberg correction (q < 0.05), and de novo motifs were filtered against dinucleotide-scrambled sequences to exclude compositional false positives. No known motif enrichment was detected in age-independent DMRs (all p-values = 1) and all 16 de novo motifs were flagged as false positives.

### Gene Ontology Annotation of DMR Associated Genes

The 294 age-independent DMRs were mapped to proximal genes using HOMER’s annotatePeaks.pl^85^ with the default association window and GO term enrichment among those genes was evaluated using findGO.pl. GO term enrichment significance was calculated using the cumulative hypergeometric distribution. Hypomethylated non-DMR regions from the same samples served as the background model. Resulting terms were filtered by nominal p-value (p < 0.05). All analyses were performed against the human reference genome GRCh38 (hg38)^67^.

### Pyrosequencing

Genomic DNA extracted for ONT sequencing from samples (39 uRPL and 30 fertile controls) were used for pyrosequencing validation. For each sample, 500 ng of genomic DNA was bisulfite converted using the EpiTect Bisulfite Conversion Kit (Qiagen), and regions of interest were amplified using the PyroMark PCR Kit (Qiagen). The Chr1 AluY DMR locus (chr1:161235257-161235567; 311 bp) was selected for pyrosequencing validation based on a combination of statistical significance, effect size, and biological relevance: the DMR spans an entire AluY element and is annotated to the upstream regulatory region of NR1I3 (1,253 bp from the transcription start site), which encodes the constitutive androstane receptor with reported roles in human hepatogenesis during embryonic development^86^ and regulation of germ cell homeostasis in mouse models^87^. Two internal sequencing regions were assayed within this amplicon: a 5-CpG region (chr1:161235409-161235461) and an adjacent 4-CpG region (chr1:161235547-161235568), covering 9 CpG sites in total. The H19 ICR (chr11:1999924-1999996; 7 CpG sites) was selected as a comparator locus based on prior reports at this region in uRPL sperm^16,17,30^, which our ONT whole-methylome profiling did not recapitulate, enabling independent orthogonal assessment of this locus. Also, we assessed genome-wide LINE1 methylation to confirm the absence of group differences observed by ONT, to test whether hypomethylation extends beyond AluY to other repeat families, and to independently examine the LINE-1 hypomethylation previously reported in RPL sperm^16^.

The 5-CpG region was amplified by nested PCR. The first reaction used an initial denaturation at 95°C for 15 minutes, followed by 45 cycles of 94°C for 30 seconds, 64°C for 30 seconds, and 72°C for 30 seconds, with a final extension at 72°C for 10 minutes. The second reaction used an initial denaturation at 95°C for 15 minutes, followed by 35 cycles of 94°C for 30 seconds, 60°C for 30 seconds, and 72°C for 30 seconds, with a final extension at 72°C for 10 minutes. The adjacent 4-CpG region and LINE1 were amplified by standard PCR using an initial denaturation at 95°C for 15 minutes, followed by 45 cycles of 94°C for 30 seconds, 54°C for 30 seconds, and 72°C for 30 seconds, with a final extension at 72°C for 10 minutes. The H19 ICR was amplified by nested PCR. Both the first and second reactions used an initial denaturation at 95°C for 15 minutes, followed by 45 cycles of 94°C for 30 seconds, 60°C for 30 seconds, and 72°C for 30 seconds, with a final extension at 72°C for 10 minutes. PCR products (10 μL) were sequenced using PyroMark Advanced reagents on a PyroMark Advanced Q24 instrument and analyzed using PyroMark Q24 Advanced software (Qiagen). Pyrosequencing primers were designed using PyroMark Assay Design software, except for the H19 ICR assay for which previously published primers were used^88,89^. All primer sequences are provided in **Supplementary Table 2**.

## Data Availability

The raw sequencing data that support the findings of this study have been deposited in NCBI SRA with the Bioproject number PRJNA1534023.

## Acknowledgments

This research was funded by the National Institutes of Health, United States of America (grant# RO1HD108124-02). We thank Martin Goros for assistance with summarizing participant characteristics. We thank the REI physicians (Drs. Randal Robinson, Belinda Yauger, Sheena Rippentrop, Robert Schenken) at UT Health San Antonio for patient recruitment. We thank Dr. Samantha Schon at the University of Michigan for providing samples through their sperm biorepository.

## Author contributions

W.M. conceived and designed the study. E.K. performed sperm sample processing and DNA extraction. J.M. performed data preparation, DMR calling, and genomic distribution analysis with help from A.T. and N.S. M.K. performed the ONT sequencing with input from T.J. N.B. performed pyrosequencing under the supervision of M.S. All sequencing and bioinformatic analysis strategies were developed and performed with input from T.J., J.H., and W.M. E.K. and W.M. conceived the data visualization strategy and drafted the first version of the manuscript. All other authors provided feedback and approved the final version of the manuscript.

## Competing Interest

T.J. and J.H. declare equity interests in Renew Biotechnologies. The remaining authors declare no competing interests.

---

**Supplementary Data 1.** Genomic location of Differentially Methylated Regions

**Supplementary Data 2.** Transcription factor binding motif analysis at DMR loci

**Supplementary Data 3.** Gene Ontology-Biological Process enrichment results for DMR-associated genes

## Notes

### Competing Interest Statement

Dr. Tim Jenkins and Jonathon Hill hold equity in Renew Biotechnologies

