## Supplemental Table 1 for "Sperm AluY Epimutations in Male Partners of Couples with Unexplained Recurrent Pregnancy Loss"

**Supplementary Table 1: Prior reported roles of 14 genes involved in the 10 highly enriched biological process**

| **Gene** | **GO terms driven (refer Suppl. Data.2)** | **Reported evidence in embryo development, implantation and placentation** | **Key references** | **AluY or Non AluY DMR** |
| --- | --- | --- | --- | --- |
| SLIT2 | 1, 2, 3, 4 | Altered SLIT2/ROBO1 signaling linked to shallow trophoblast invasion and placental angiogenesis inhibition in RPL decidua. | Chen et al. 2021; Li et al. 2017 | Hypo- AluY DMR |
| NRP1 | 1, 2, 3, 5, 6 | VEGF co-receptor expressed in human decidua, villi and invading cytotrophoblast across all trimesters and linked to embryonic implantation and placentation. | Baston-Buest et al. 2011 | Hypo-DMR;  Non-AluY |
| SEMA3A | 1, 2, 3 | NRP1-SEMA3A promotes vascularization at embryo-maternal interface during the implantation window | Zhang et al. 2026 | Hypo- AluY DMR |
| ALCAM | 1, 2, 3 | Expressed on human blastocysts and endometrial epithelial cells in developmentally stage-specific manner; homophilic ALCAM-ALCAM adhesion proposed as initial embryo-endometrium recognition mechanism | Fujiwara et al. 2003 | Hypo- AluY DMR |
| EPHA4 | 3, 4, 5, 6, 8, 9 | Eph receptor family plays a role in trophoblast migration and placental angiogenesis; EphA4 detected in endometrium during implantation; Eph-Ephrin system regulates blastocyst attachment and spreading | Fu et al., 2012; Goldman-Wohl et al. 2004 | Hypo- AluY DMR |
| EPHA3 | 5, 6, 8 | Ephrin-A3 validated as functional target of miR-210 regulating cell migration and vascular remodelling; originally named "Human Embryo Kinase" reflecting embryonic discovery context | Goldman-Wohl et al. 2004; Zhang et al., 2012 | Hypo-DMR;  Non-AluY |
| JAG1 | 7 | Expressed in decidual vasculature and trophoblasts during placentation; specifically reduced in invasive trophoblasts from women with preeclampsia | Levin et al. 2017 | Hyper-DMR;  Non-AluY |
| SOX2 | 7 | Essential for trophectoderm formation in preimplantation embryo; Sox2-null embryos fail to form trophectoderm and die after implantation | Keramari et al. 2010 | Hypo- AluY DMR |
| CDH2 | 7, 9, 10 | N-cadherin promotes placental trophoblast invasion; BMP2-mediated CDH2 upregulation enhances extravillous trophoblast invasive capacity | Zhao et al. 2018 | Hyper- AluY DMR |
| CDH11 | 4, 9, 10 | Hormonally regulated marker of endometrial stromal decidualization; expression increases under progesterone alongside biochemical markers of decidualization | Chen et al. 1999 | Hypo- AluY DMR |
| FUT10 | 7 | FUT10 embryo-specific splice variants were expressed in human embryos | Mollicone et al. 2009 | Hypo- AluY DMR |
| CDH8 | 9, 10 | Essential for cortical neuron development in embryo. | Memi et al. 2019 | Hypo- AluY DMR |
| CDH13 | 9, 10 | Placental CDH13 was hypermethylated in fetal overgrowth | Yang et al., 2022 | Hypo- AluY DMR |
| TNFRSF21 | 5, 6 | Embryonic brain vascular development | Tam et al., 2012 | Hypo-DMR;  Non-AluY |

**Supplementary Table 2: Primers used for the pyrosequencing validation**

| **Region** | **Primers** | **Target location (hg38)** | **Amplicon length** |
| --- | --- | --- | --- |
| H19 DMR | Outer Forward: 5′-TTTTTGGTAGGTATAGAGTT-3′ | chr11:1999924-1999996 | 231 |
|  | Outer Reverse: 5′-AAACCATAACACTAAAACCC-3′ |  |  |
|  | Nested Forward: 5′-TGTATAGTATATGGGTATTTTTGGAGGTTT-3′ |  |  |
|  | Nested Reverse*: 5′-TCCTATAAATATCCTATTCCCAAATAACC-3′ |  |  |
|  | Sequencing: 5'-TGGTTGTAGTTGTGGAAT-3' |  |  |
| AluY (Region 1) | Forward: 5'-GGGGTTTTATTGTGTTAGTTAGAA-3' | chr1:161235257-161235767 | 146 |
|  | Reverse*: 5'-AACCACACTAACTTCACTATCAT-3' |  |  |
|  | Sequencing: 5'-GTTTTTTAAAGTGTTGGGA-3' |  |  |
| AluY (Region 2) | Outer Forward: 5′-CCATGTGGCTATCTAGGAGAAAAGCATTTC -3′ | chr1:161235409-161235461 | 186 |
|  | Outer Reverse: 5′-GCCACACTGACTTCACTGTCATTCTTAGA -3′ |  |  |
|  | Nested Forward: 5′-TTATTTAGGTTGGAGTGTAGTGG-3′ |  |  |
|  | Nested Reverse*: 5′-AACCATTCTAACTAACACAATAAAAC-3′ |  |  |
|  | Sequencing: 5'-AGTAGTTGGGATTATAGG-3' |  |  |
| LINE 1 | Forward: 5'-GGGGAGGAGTTAAGATGG-3'  Reverse*: 5'-CACTATCTAACACTCCCTAATAAAAT-3'  Sequencing: 5'-GAGGAGTTAAGATGGT-3' | All chromosomes (GGGGGAGGAGCCAAGATGGCCGAATAGGAACAGCTCCGGTCTACAGCTCCCAGCGTGAGCGACGCAGAAGACGGG) | 134 |
