## Supplemental Figures 1,2,3 for "Sperm AluY Epimutations in Male Partners of Couples with Unexplained Recurrent Pregnancy Loss"

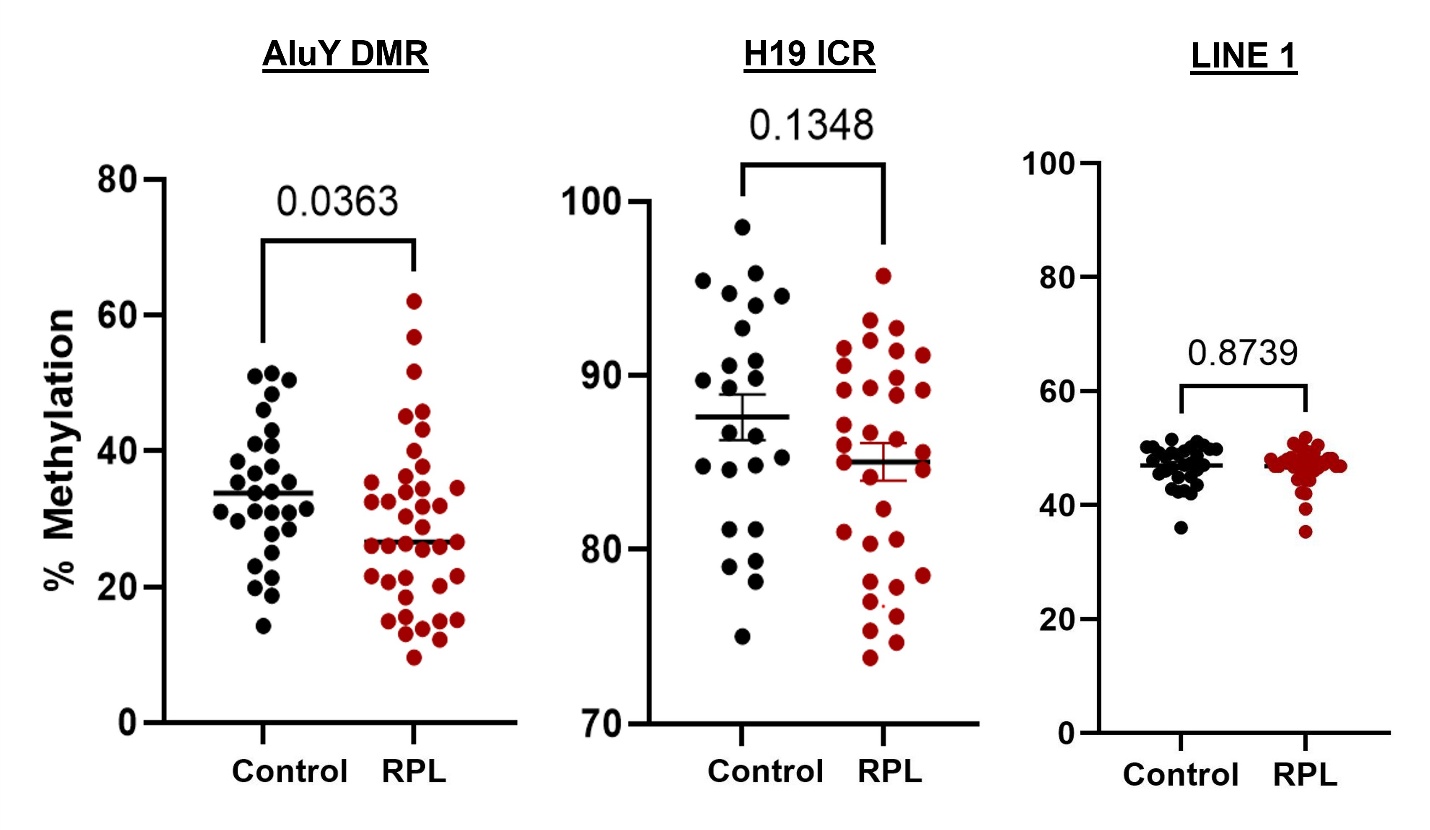

Supplementary Figure 1: Pyrosequencing validation of AluY DMR, H19 ICR and LINE1 methylation. a, Mean bisulfite pyrosequencing methylation at a representative AluY DMR (chr1:161,235,257-161,235,567; 9 CpG sites across two sequencing regions), differed significantly (p = 0.03) between fertile donors and uRPL patients. b, Mean bisulfite pyrosequencing methylation at the H19 imprinting control region (chr11:1,999,924-1,999,996; 7 CpG sites), showing no significant difference between groups (p=0.13). c, Genome-wide LINE1 methylation didn’t differ significantly between groups (p=0.8). All three results are concordant with the corresponding ONT-based findings at these loci.

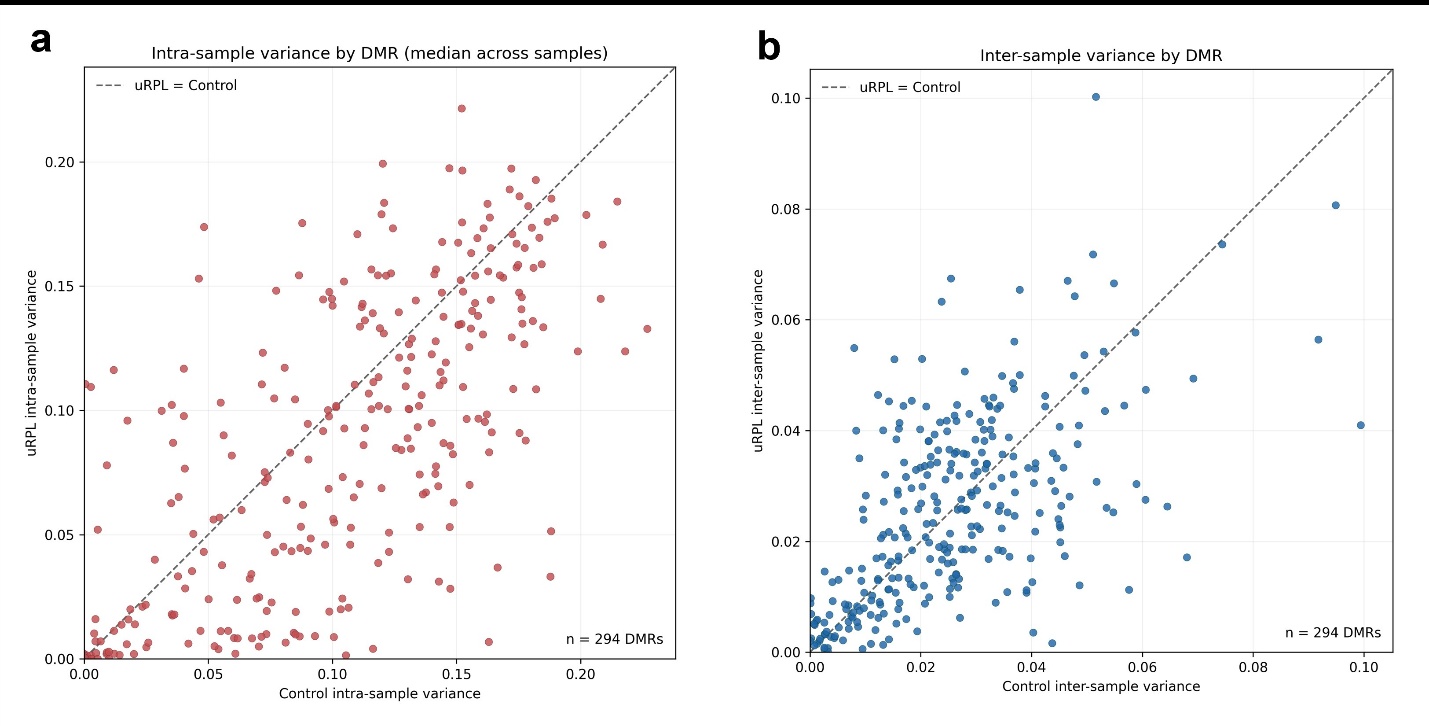

**Supplementary Figure 2: Intra- and inter-sample methylation variance at DMR loci. a,** Per-DMR scatter plot displaying intra-sample variance between fertile donors (X-axis) and uRPL patients (Y-axis), for each of the 294 DMRs. **b,** Per-DMR scatter plot displaying inter-sample variance between fertile donors and uRPL patients, for each of the 294 DMRs. Majority of DMRs exhibit comparable variance between groups.

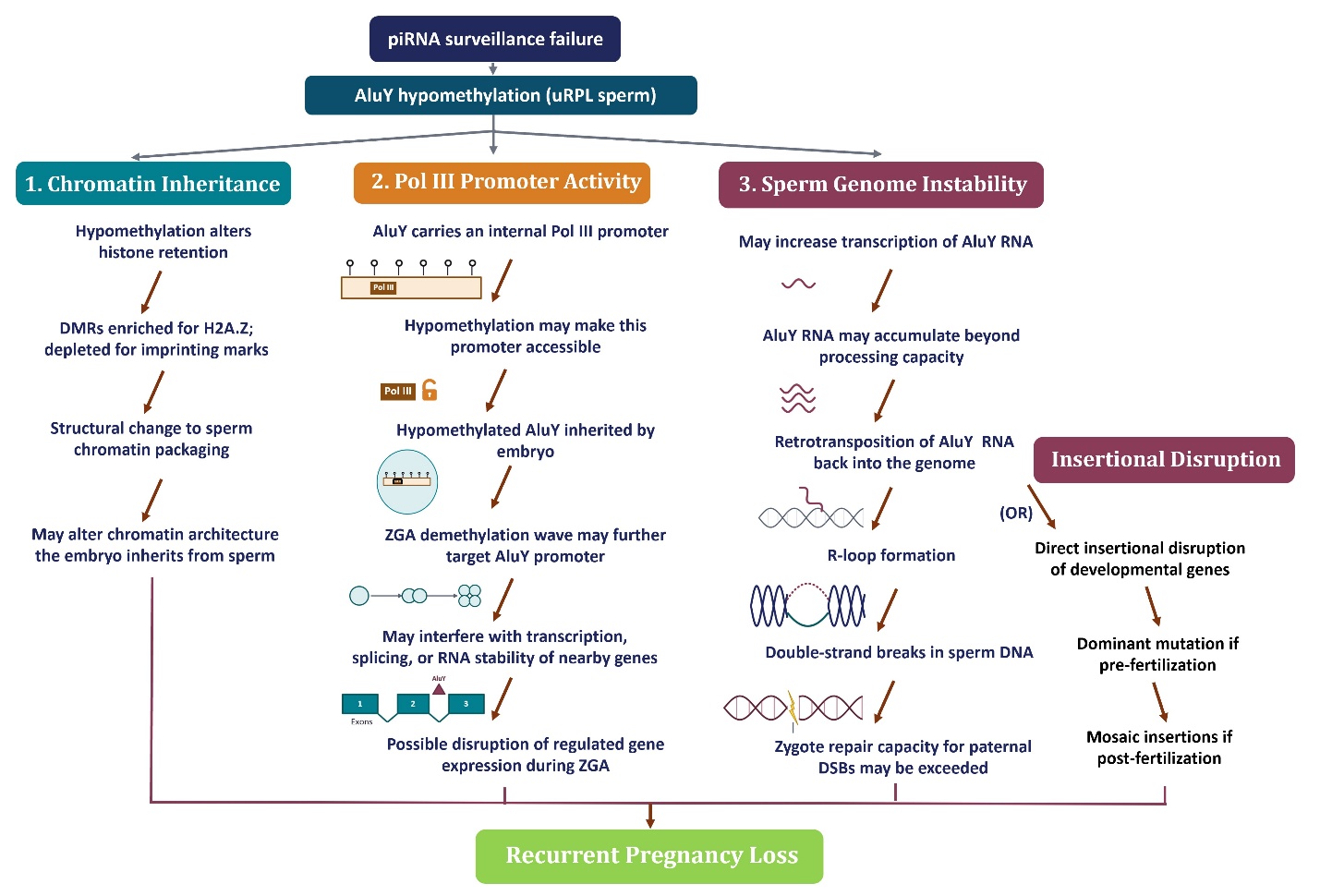

**Supplementary Figure 3:** Hypothetical mechanistic pathways linking paternal AluY hypomethylation to recurrent pregnancy loss
